# *Wolbachia* genomics reveals a potential for a nutrition-based symbiosis in blood-sucking Triatomine bugs

**DOI:** 10.1101/2022.09.06.506778

**Authors:** Jonathan Filée, Kenny Agésilas-Lequeux, Laurie Lacquehay, Jean Michel Bérenger, Lise Dupont, Vagner Mendonça, João Aristeu da Rosa, Myriam Harry

**Affiliations:** UMR Evolution, Génomes, Comportement, Ecologie (EGCE), IDEEV, Université Paris Saclay-CNRS-IRD, 12 route 128, Gif-sur-Yvette, France; Muséum National d’Histoire Naturelle, Département Systématique & Evolution (Entomologie), 45, rue Buffon, 75005 Paris, France; Aix Marseille Université, IRD, AP-HM, SSA, VITROME, IHU-Méditerranée Infection. Marseille, France; Université Paris Est Créteil, Sorbonne Université, CNRS, INRA, IRD, Université Paris-Diderot, Institut d’Ecologie et des Sciences de l’Environnement de Paris, Créteil, France; Universidade Federal do Piauí, Departamento de Parasitologia e Microbiologia/CCS, Teresina/PI, Brazil; Universidade Estadual Paulista (Unesp), Faculdade de Ciências Farmacêuticas, Rodovia Araraquara-Jaú, Araraquara, São Paulo, Brasil

**Keywords:** *Wolbachia*, *Rhodococcus rhodnii*, *Rhodnius*, blood-sucking bugs, Nutritional symbiosis, Comparative genomics, biotin operon

## Abstract

The nutritional symbiosis promoted by bacteria is a key determinant for adaptation and evolution of many insect lineages. A complex form of nutritional mutualism that arose in blood-sucking insects critically depends on diverse bacterial symbionts that supplement the diet of their nutrient-poor hosts with B vitamins. For instance, the triatomine bug *Rhodnius prolixus*, one of the main vectors of the Chagas disease in humans, is known to maintain a nutritional symbiosis with the gut symbionts *Rhodococcus rhodnii*.

In this study, we show that *Wolbachia* symbionts are also widely distributed in the *Rhodnius* genus. We have screened a large set of *Rhodnius* blood-sucking bugs samples belonging to 17 different species and to the three phylogenetic groups, *prolixus, pallescens* and *pictipes*. We assembled 13 genomes of *Wolbachia* infecting eight *Rhodnius* species from *prolixus* and *pictipes* groups. We demonstrate that these *Wolbachia* belong to supergroup F and are closely related to *Wolbachia* infecting the bedbug *Cimex lectularius* (*w*Cle). Although bedbugs and triatomines are very distantly related hemipteran bugs, the genomes of their respective *Wolbachia* were highly similar, suggesting recent horizontal host switches. We also show that *Rhodnius Wolbachia* genomes infecting the *prolixus* group encode intact biotin operon, the hallmark of nutritional symbiosis in bedbugs. This operon is lacking from all the other *Wolbachia* infecting *R. pictipes*. Finally, host genome analyses provide evidence of massive *Wolbachia*-to-*Rhodnius* gene transfers in almost samples, providing footprints of past infections that support a widespread and probably ancient symbiotic association between *Wolbachia* and triatomine bugs.

Our results suggest that both *Wolbachia* and *R. rhodnii* gut symbionts and their *Rhodnius* host maintain a highly prevalent symbiotic relationship, in which the vertically-inherited *Wolbachia* has the metabolic potantial to ensure or complement, the nutritional mutualism provided by the gut symbionts. Specific loss of the biotin operon in some symbiont genomes suggests that the boundaries between obligatory mutualism, facultative mutualism and parasitism in *Wolbachia* are transient and fluid, supporting a dynamic process of transition and reversion from one state to another.

## Introduction

The triatomine bugs (Hemiptera, Reduviidae, Triatominae) are blood-sucking vectors of *Trypanosoma cruzi*, the etiological agent of the Chagas disease that affects about 6 million people in Latin America (PAHO, 2020). The most famous triatomine bug, *Rhodnius prolixus*, has served as a model insect particularly for the study of physiological phenomenons. It was suggested by Wigglesworth that this triatomine bug critically depends on gut symbiotic bacteria for larvae development (Wigglesworth 1936). Later, Baines proposed that a bacterial symbiont, *Rhodococcus* (former *Nocardia) rhodnii*, living in the midgut of the bugs, provides their hosts with B-group vitamins such as biotin, nicotinamin, thiamin, pyridoxin, or riboflavin (Baines 1956). This phenomenon, called “nutritional mutualism”, is widespread in insects and involves a large variety of microbes and metabolic capabilities (Sudakaran et al. 2017). *Rhodococcus* symbionts live in the midgut of the bugs and are orally acquiring, ingested by newly hatched nymphs, since they are transmitted maternally through the smearing egg-surface with symbiont-containing feces contamination or vertically via coprophagy of symbiont-containing adult feces (Wigglesworth 1936). The phylogenetic distribution of *Rhodococcus* among the *Rhodnius* genus is unknown, as they have only been isolated in *R. prolixus* and *R. ecuadoriensis* (Rodríguez et al. 2011). The most puzzling aspect of this symbiotic relationship is the true nature of the metabolic benefits provided by *R. rhodnii*. Many contradictory results tend to demonstrate that the nutritional mutualism between *R. rhodnii* and *Rhodnius* is not strictly obligatory but depends mostly on rearing condition, host bloods or symbiont strains. In some studies, the larvae development of symbiont-free insects is fully restored using blood supplemented with B vitamins (Lake and Friend 1968; Auden 1974). But in others, the requirement of this supplementation depends on the type of blood diets: *Rhodnius* bugs fed on mouse blood do not require B-vitamin supplementation, whereas supplementation is mandatory for bugs fed on rabbit blood (Baines 1956; Nyirady 1973). Moreover, bug development was similar using the *Rhodococcus* wild strain or auxotrophic mutants that are not able to produce B vitamins, indicating that the *Rhodococcus/Rhodnius* symbiosis may imply other kinds of metabolic benefits (Hill et al. 1976). This latter study also opens the possibility that secondary symbionts may reinforce or rescue the mutualistic relationships (Hill et al. 1976). This role could be played by bacteria belonging to the genus *Wolbachia*.

*Wolbachia* are endosymbiotic, maternally inherited bacteria that infect a large array of arthropods and nematodes (Werren et al. 2008). In most cases, *Wolbachia* are facultative symbionts that manipulate their hosts to increase their own transmission, causing negative fitness consequences (Werren et al. 2008). However, in insects, exceptions exist in which *Wolbachia* have established obligatory symbiosis. For example, in parasitoid wasps the removing symbiotic *Wolbachia* bacteria specifically inhibits oogenesis (Dedeine et al. 2001). But also, *Wolbachia*-free insects exhibit deficiencies in growth and fecundity but are rescued by a blood diet supplemented with biotin (Hosokawa et al. 2010). Indeed, in the bedbug *Cimex lectularius*, it was demonstrated that *Wolbachia* infecting *Cimex* hosts (*w*Cle) inhabit specialized organs, the bacteriomes, located close to the gonads (Hosokawa et al. 2010). Genomic and biochemical studies have shown that the *w*Cle genome encoded a functional biotin operon that has been acquired via a lateral gene transfer from a co-infecting symbiont (Nikoh et al. 2014). If biotin appears to be the cornerstone of the B-vitamin nutritional mutualism, riboflavin provisioning by *w*Cle might also play a metabolic role in the association (Moriyama et al. 2015). *Wolbachia* was also found in termite bacteriocytes (wCtub), this localization suggesting that these symbionts may be nutritional mutualists especially for *Cavitermes tuberosus* as *w*Ctub harbours the *bioA* gene involved in the biotin (vitamin B7) synthesis pathway (Hellemans et al. 2018). *Wolbachia* symbiotic role was also described in the planthoppers, *Laodelphax striatellus* and *Nilaparvata lugens*, the presence of *Wolbachia* rescuing insect fecundity deficit, and the genomic analysis showing that *Wolbachia* strains (*w*Lug, *w*StriCN) from these two planthoppers encoded complete biosynthesis operons for biotin and riboflavin (Ju et al. 2020). The biotin operon was also found in two *Nomada* bees, *N. flava* (*w*Nfa) and *N. leucophthalma* (*w*Nleu), but as these bees feed on pollen which typically contains high levels of B vitamins, a *Wolbachia* role in nutritional provisioning seems unlikely for these species (Gerth and Bleidon 2017).

Phylogenetically, *Wolbachia* have been clustered in at least 13 lineages, denominated “supergroups” named A-F, H-Q, and S (Kaur et al. 2021). The biotin operon (bioABCDFH loci) has been identified in three *Wolbachia* supergroups A, B and F. It was shown that *w*Nleu and *w*Nfla belong to the supergroup A (Gerth and Bleidon, 2017), *w*Lug and *w*StriCN to the supergroup B (Ju et al. 2020), and wCle to the supergroup F (Gerth et al. 2014; Nikoh et al. 2014). Supergroups A and B were described first and are most commonly found among arthropod species but also the supergroup F, found in a large diversity of insects, such as bedbugs, termites, bush crickets, louse-flies, weevils, cockroaches or ant lions (Ros et al. 2009; Kaur et al. 2021). In Triatominae, although *Wolbachia* have been identified in *R. pallescens* based on 16S signatures (Espino et al. 2009) and in the *R. prolixus* genome (Mesquita et al. 2015), nothing is known about the origin, distribution or type of symbiotic relationships between the blood-sucking bugs and their *Wolbachia*.

In this study, we first tested the respective prevalence of *Wolbachia* and *R. rhodnii* in a large sampling of *Rhodnius* belonging to sixteen different species. The *Rhodnius* genus comprises twenty-four species including twenty-one *Rhodnius* species and three *ex-Psammolestes* species as demonstrated by phylogenomics studies and is divided into three major groups, *pictipes*, *prolixus* and *pallescens* (Filée et al., 2022). In order to understand the evolutionary origin and the possible metabolic roles of the *Wolbachia*-infecting *Rhodnius* species, we sequenced and analysed the *Wolbachia* genomes. Finally, we also examined the *Rhodnius* genomes for the presence of gene transfers from *Wolbachia* with the aim of using them as the footprints of past infections. Albeit the gut *R. rhodnii* symbionts play the role of the obligatory nutritional symbiont by providing B-vitamins to the *Rhodnius* triatomine, our findings support that *Wolbachia* could potentially provide a benefit to their hosts to ensure or to complement the nutritional mutualism provided by the gut symbionts.

## Materials and methods

### Insect sampling and preparation

We used 117 females of *Rhodnius* specimens dispatched in 39 populations and belonging to 17 species (Table 1). Among them, 70 females (14 species) originated from the field and 47 females (9 species) were reared at the Brazilian Insetário de Triatominae da Faculdade de Ciências Farmacêuticas, Universidade Estadual Paulista (UNESP). All these 117 specimens were screened for *Wolbachia* infection by PCR experiments after DNA extraction from genital tissues using the Qiagen DNEasy tissue kit. A subsample of 36 specimens including *Wolbachia*-free and infected insects were used for high throughput DNA sequencing after DNA extraction from legs and alary muscles. For the 36 specimens used for genomics, the *Rhodoccocus* presence/absence analysis was also performed using DNA extracted from gut and internal organs. Species determination was performed using a phylogenomic study (Filée et al., 2022) using both mitochondrial (13 protein-coding mitochondrial genes) and nuclear data (rDNA, 51 protein-coding nuclear genes). Few species showed mitochondrial introgression between close relative species of the same phylogenetic group (Table 1) which do not change the overall topology of the tree nor the interpretation of the distribution pattern of the symbionts.

**Table 1:** Main characteristics of the *Rhodnius* populations/strains analyzed in this study. Species are listed in the same order than the phylogeny (Figure 1). *Rhodnius* groups are indicated as Pall: *pallescens*, Pic: *pictipes*, Pro: *prolixus*. Mitochondrial introgression is indicated. Presence of *Wolbachia* infection is stated with indications regarding the number of individuals tested using PCR experiments with *coxA* and *ftsZ* primers and the number of infected specimens. Infected species are indicated in bold.

| Samples | Group | Species | Origin | Field/Lab | Nb<br>*Wolbachia*<br>infected | Total<br>tested |
| --- | --- | --- | --- | --- | --- | --- |
| ECUE | Pall | *R. ecuadorensis* | Peru | F | 0 | 1 |
| ECUD | Pall | *R. ecuadorensis* | PD, Colombia | F | 0 | 1 |
| R1J | Pall | *R. pallescens* | Colombia | L (CTA s/R ) | 0 | 5 |
| GS | Pall | ***R. pallescens*** | Galevar Sucre, Colombia | F | 1 | 4 |
| VA | Pall | ***R. pallescens*** | Vegachi Antioqua, Colombia | F | 4 | 8 |
| SO | Pall | ***R. pallescens*** | San Onofre, Colombia | F | 5 | 10 |
| R10B | Pall | *R. colombiensis* | Tolima, Colombia | L (CTA 050) | 0 | 5 |
| U-StaWY | Pic | ***R. stali*** | Alto Beni, Bolivia | F | 2 | 2 |
| V-StaWZ | Pic | ***R. stali*** | Bolivia | F | 3 | 3 |
| R9WX | Pic | ***R. brethesi*** | Igarapé Tucunaré, Brazil | L (CTA 222) | 5 | 5 |
| BRE25W<br>B | Pic | ***R. brethesi*** | Amazonia, Brazil | F | 1 | 1 |
| R6WV | Pic | ***R. pictipes*** | Belem, PA, Brazil | L (CTA 072) | 4 | 5 |
| PIC2WA | Pic | ***R. pictipes*** | Para, Brazil | F | 1 | 5 |
| R63K | Pic | *R. pictipes* | Belem, Para, Brazil | L (CTA 072) | 0 | 1 |
| PIC34W<br>C | Pic | ***R. pictipes*** | Belem, Brazil | F | 1 | 1 |
| PIC3L | Pic | ***R. pictipes*** | Acarouany, French Guyane | F | 4 | 8 |
| Ama1A | Pic | ***R. amazonicus*** | Bélizon, French Guyana | F | 1 | 1 |
| MILEP | Pro | *R. prolixus* | Para, Brazil | F | 0 | 1 |
| RobQ | Pro | ***R. prolixus*** | Para, Brazil | F | 1 | 3 |
| ProN | Pro | *R. prolixus* | Colombia | L (CTA 080) | 0 | 3 |
| ProM | Pro | *R. prolixus* | Colombia | L (CTA 077) | 0 | 3 |
| Pro10O | Pro | *R. prolixus* | Estado Guarico, Venezuela | F | 0 | 1 |
| NEGP | Pro | *R. prolixus* | Piaui, Para, Brazil | F | 0 | 0 |
| R8F | Pro | *R. montenegrensis* | Montenegro, RO, Brazil | L (CTA 087) | 0 | 5 |
| R4H | Pro | *R. prolixus* (mt *R. montenegrensis*) | Brazil | F | 0 | 1 |
| RobR | Pro | *R. prolixus* (mt *R. montenegrensis*) | Peru | F | 0 | 2 |
| ROBB | Pro | *R. marabaensis* | Maraba, Brazil | F | 0 | 0 |
| ProYRP | Pro | *R. prolixus* (mt *R. robustus*) | Cojedes, Venezuela | F | 0 | 3 |
| Rob5s | Pro | *R. robustus* | French Guyana | F | 0 | 1 |
| R3WT | Pro | ***R. neglectus*** | Taquaruçu, MS, Brazil | L (CTA) | 5 | 5 |
| 5WU | Pro | ***R. neglectus*** | Frutal, MG, Brazil | L (CTA 061) | 5 | 5 |
| NASP | Pro | *R. nasutus* | Piaui, Para, Brazil | F | 0 | 0 |
| MILE | Pro | *R. milesi* | Brazil | F | 0 | 1 |
| INCP | Pro | *R. neglectus* (mt *R. nasutus*) | Piaui, Para, Brazil | F | 0 | 0 |
| R7WU | Pro | ***R. nasutus*** | Brazil | L (CTA 054) | 5 | 5 |
| NasG | Pro | *R. nasutus* | Piaui, Brazil | F | 0 | 1 |
| PSAM | Pro | *R. (Psammolestes) tertius* | Brazil | F | 0 | 1 |
| NEII | Pro | *R. neivai* | Maracay, Venezuela | F | 0 | 3 |
| DomC | Pro | *R. domesticus* | Santa Catalina, Brazil | F | 0 | 3 |
| Field |  | 14 species **(6)** |  |  | 23 (35%) | 66 |
| Strains |  | 9 species **(4)** |  |  | 24 (51%) | 47 |
| Total |  | 17 species **(8)** |  |  | 47 (42 %) | 113 |

### PCR experiments

The presence of *Wolbachia* was detected using standard *coxA* and *ftsZ* primers in the 120 samples as previously described (Werren and Windsor 2000). We designed specific *Rhodococcus rhodnii* primers targeting the *16s* and the *groEL* genes using the Primer-BLAST software (Ye et al. 2012) seeded with corresponding sequences derived from the *R. rhodnii* LMG5362 whole genome sequence (Genbank access: GCA_000389715). *16S* primers correspond to 5’ ACATGCAAGTCGAGCGGTAA (forward) and 5’ GTGTCTCAGTCCCAGTGTGG (reverse) and *groEL* to 5’ GTGGTCTCGTCCTTGGTGAC (forward) and 5’ CTGCTCTACCGCGACAAGAT (reverse). Standard PCR reaction mixtures contained deoxynucleoside triphosphates (10μM, 0.2 μl per tube), both primers (10μM, 1 μl each per tube), Go *Taq* flexi polymerase from Promega (5U/μl, 0.1μl per tube), MgCl2 (50mM, 3 μl per tube), 1X buffer (5 μl per tube), a DNA sample (1 μl per tube) and water to reach a final volume of 25μl (13.7 μl). PCR products were then sequenced at the Eurofins Scientific PlateSeq service (Moissy-Cramayel, France). PCR sequences were then searched with BLAST (Altschul et al. 1990) against a NR database to verify that the PCR products correspond to *Wolbachia* and *R. rhodnii* sequences using the followed criteria: first BLAST hit and sequence identity >90% for *Wolbachia* for which we do not have any know sequence infecting *Rhodnius* species and >99% for *R. rhodnii* for which a reference genome is available.

### *Rhodnius* and *Wolbachia* genome sequencing and assembly

DNA samples of 36 triatomines (550-7000μg) were subjected to whole-genome shotgun sequencing using Illumina HiSeq, corresponding to a total of 15 to 25 Gb data per sample (100 bp paired-end, Imagif platform, Gif-sur-Yvette, France). Assembly was carried out with the SOAPdenovo2 software (Luo et al. 2012) with k-mers estimated using the KmerGenie program (Chikhi and Medvedev 2014). As *Wolbachia* genes are frequently inserted into the host genomes, it is important to filter the sequences to keep the contigs that align with *Wolbachia* sequences present in the sequence database with the exclusion of sequences that also match the reference *R. prolixus* genome. To reach this goal, we used the approach described in Kumar and Blaxter 2011 (Kumar and Blaxter 2011): All the contigs of the assemblies were first searched with BLASTN (Altschul et al. 1990) against a non-redundant Genbank database from the National Center for Biotechnological Information (NCBI). Contigs were assigned to *Wolbachia* using the following criteria: (1) first match with *Wolbachia* sequences using 1e-20 e-value cut-off, (2) align on at least 50% of its length on *Wolbachia* sequences, and (3) do not match with the *Wolbachia* sequences integrated into the *R. prolixus* C3 reference genome (available at https://www.vectorbase.org/), therefore these sequences were masked (replaced by “N”). Contigs were finally discriminated using their read coverage using the following method: for a given species, raw reads were mapped using BWA (Li and Durbin 2010) to the corresponding species assembly to compare the level of coverage between the putative *Wolbachia* contigs and the remaining contigs.

### *Wolbachia* and *Rhodococcus* genome analysis

Study of genome conservation among *Wolbachia* genomes was carried out using the BRIG software (Alikhan et al. 2011) and whole genome alignments using Vista (Poliakov et al. 2014) with default parameters. Identification of Insertion Sequences (IS) was carried out by querying the IS finder database (Siguier et al. 2006). Analyses of the B-vitamin genes were conducted with TBLASTN (Altschul et al. 1990) searches seeded with the *Escherichia coli* and the *Cimex Wolbachia* homologs of each gene. Analysis of the B-vitamin genes in four complete genomes of the triatomine gut symbionts belonging to the genera *Rhodococcus* was also conducted in a similar way (accession numbers: APMY00000000.1, *Rhodococcus rhodnii* LMG 5362; BCXD00000000.1, *Rhodococcus rhodnii* NBRC 100604; FNDN00000000.1, *Rhodococcus triatomae* strain DSM 44892; AODO00000000.1, *Rhodococcus triatomae* BKS 15-14). Orthologs were aligned using MAFFT (Katoh et al. 2002) and carefully checked for deletions and/or stop-codons. The rate of synonymous and non-synonymous substitutions was computed using the KaKs_Calculator2.0 package using the Model Selection (MS) option (Zhang et al. 2006).

Moreover, we checked if *Wolbachia* genes were laterally transferred into the *Rhodnius* genomes using two criteria (Chung et al. 2017). *Wolbachia* genes were considered as inserted into the host genome only if their 3’ and 5’ flanking sequences (both ends) align to the reference *R. prolixus* genome. Thus, we first aligned the raw reads against the *Rhodnius Wolbachia* genomes using BWA. The aligned reads were then assembled using Trinity (Grabherr et al. 2011) and the assembled sequences were aligned on: (1) the masked *Rhodnius prolixus* C3 genome using 1e-20 e-value cutoff, retaining the sequences >100nt that align both on the 5’ and the 3’ ends and (2) the *Rhodnius Wolbachia* genome assemblies with a nucleotide similarity threshold of 95%. Putative cases of lateral gene transfers were then visualized using the IGV genome browser by mapping them on the *w*Cle genome (Robinson et al. 2011). We also used the level of read coverage to distinguish the contigs deriving from the symbionts to those hosted by the genome as proposed by Kumar and Blaxter (Kumar and Blaxter 2011). In addition, we analysed the lateral *Wolbachia* gene transfers in the published *R. prolixus* reference genome, using a nucleotide identity threshold of 90% as a criterion.

### Phylogenetic analysis

*Rhodnius* phylogeny was carried out using an alignment of the mitochondrial genomes (Filée et al., 2022). *Wolbachia* phylogeny was carried out using the genomes of 12 *Wolbachia* strains representing the supergroups A, B, C and D, in addition to 4 outgroups retrieved from the dataset of 90 conserved genes identified previously (Comandatore et al. 2013). For the *w*Cle and 14 *Rhodnius Wolbachia* genomes, we identified orthologs of these conserved genes using reciprocal BLASTN searches. As genes with incomplete taxon sampling were excluded, the final data set comprised 80 ortholog genes for a total of 23,700 nucleotides. A similar approach was conducted for the biotin phylogeny, using the amino acid dataset published elsewhere (Gerth and Bleidorn 2016).

Alignments were performed with MAFFT (Katoh et al. 2002) and visualized with Aliview (Larsson 2014). Maximum Likelihood phylogenies (ML) were reconstructed with PhyML (Guindon and Gascuel 2003), using the best-fit nucleotide substitution model GTR for *Wolbachia* phylogeny and the JTT, best-fitted model of protein evolution, for the biotin phylogeny. The reliability of branching patterns in ML trees was assessed with 1,000 non-parametric bootstraps.

## Results

### Distribution of *Wolbachia* and *Rhodococcus rhodnii* in the genus *Rhodnius*

The results of the PCR screening revealed a different pattern of presence/absence in *Wolbachia* and *R. rhodnii* (Figure 1). Out of the 113 specimens targeted by PCR using specific *Wolbachia coxA* and *ftsZ* primers, 42% were *Wolbachia*-infected, corresponding to 8 species, namely *R. amazonicus, R. brethesi, R. nasutus, R. neglectus, R. pallescens, R. pictipes, R. prolixus*, and *R. stali* (Table 1 and Figure 1). The infection was differentiated according to the origin of the specimens. Infection by *Wolbachia* was observed in 51% (corresponding to 4 different species, namely *R. brethesi, R. nasutus, R. neglectus*, and *R. pictipes*) of the specimens reared in the laboratory (strains) and in only 3% (corresponding to 6 different species namely *R. amazonicus, R. brethesi, R. pallescens, R. pictipes, R. prolixus*, and *R. stali*) of the specimens from the field. It is noteworthy that for a given population/strain all the specimens were not always infected. For example, for the *R. pictipes* Pic3L population, *Wolbachia* were detected in 4 out of 8 specimens. On the other hand, *R. rhodnii* was present in 100% of the tested samples whatever the *Wolbachia* infection status (Figure 1). *Wolbachia coxA* and *ftsZ* sequences displayed high levels of similarity with the supergroup F *Wolbachia* infecting various insects (>99% nucleotide identity). Moreover, *16S* and *groEL R. rhodnii* fragments were identical to the corresponding *R. rhodnii* sequences isolated from *R. prolixus*.

**Figure 1:**
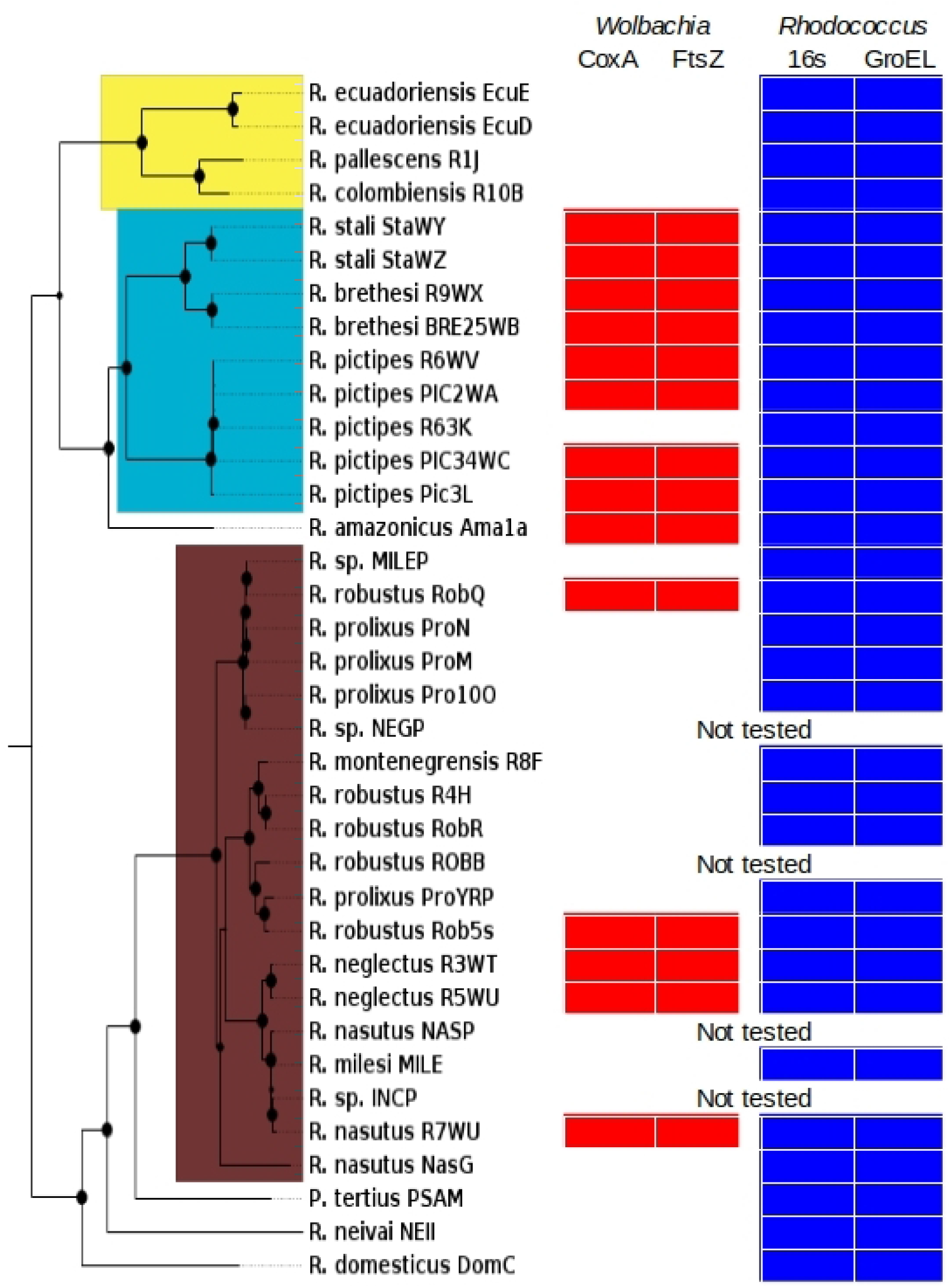
Distribution of the *Wolbachia* and the *Rhodococcus rhodnii* symbionts in the genus *Rhodnius*. Presence/absence of specific PCR products targeting *Wolbachia coxA* and *ftsZ* genes (red rectangles) and *R. rhodnii 16S* and *groEL* genes (blue rectangles) mapped on the *Rhodnius* whole mitochondrial genome maximum-likelihood phylogeny. Yellow, blue and brown groups in the tree refer to the *pallescens, pictipes* and *prolixus* groups respectively. Black circles in the phylogeny indicate the support values of each node: large circles for bootstraps >99%, small ones for supports between 90% and 99%.

### Whole genome sequence and analysis of the *Wolbachia* infecting *Rhodnius*

The whole genome of the 36 specimens of *Rhodnius* previously screened by PCR were sequenced. After genomic assemblies, contigs were assigned to *Wolbachia* if they aligned to known *Wolbachia* sequences using stringent criteria (first match with *Wolbachia* sequences using 1e-20 e-value cut-off and align on at least 50% of its length on *Wolbachia* sequences) discarding contigs that also match the *R. prolixus* reference genome to avoid *Wolbachia* sequences integrated in the host genomes. Except for *R. pallescens* samples not subjected to whole-genome shotgun sequencing, the other 14 samples tested positive by PCR for *Wolbachia* led to the reconstruction of a draft *Wolbachia* genome (Table 2). The *w*RobQ genome assembly appears to be incomplete due to the small size of the assembly, while the remaining 13 genomes have a genome size from 0.96 to 1.15 Mb (1.08 ± 0.06 Mb).

**Table 2:** Main characteristics of the *Rhodnius*-associated *Wolbachia* genome assemblies. Details include the geographic origins, genome assembly sizes and N50, the number of Insertion Sequences and the GC%.

| Samples | Species | Origin | Size (nt) | N50 (nt) | IS(#) | GC (%) |
| --- | --- | --- | --- | --- | --- | --- |
| ***prolixus* group** |  |  |  |  |  |  |
| wINCP | *R. neglectus* (mt *R. nasutus*) | Piauí, Para, Brazil | 982754 | 2399 | 24 | 36 |
| wR3WT | *R. neglectus* | Taquaraçu, Mato-Grosso do Sul, Brazil | 1158544 | 5245 | 36 | 36 |
| wR5WU | *R. neglectus* | Frutal, Minas Gerais, Brazil | 1151075 | 6061 | 36 | 36 |
| wR7WU | *R. nasutus* | Brazil | 1092395 | 4706 | 41 | 36 |
| wRob5s | *R. robustus* | French Guyana | 1090475 | 2418 | 21 | 36 |
| wROBB | *R. marabaensis* | Marabá, Brazil | 1020284 | 1891 | 30 | 36 |
| wRobQ | *R. prolixus* | Para, Brazil | 227045 | 2937 | 21 | 35 |
| ***pictipes* group** |  |  |  |  |  |  |
| wAma1A | *R. amazonicus* | Bélizon, French Guyana | 962616 | 3071 | 20 | 36 |
| wR9WX | *R. brethesi* | Igarapé Tucunaré, Brazil | 1132878 | 5051 | 28 | 36 |
| wBRE25WB | *R. brethesi* | Amazonia, Brazil | 1139249 | 7047 | 21 | 36 |
| wPIC3L | *R. pictipes* | Acarouany, French Guyana | 1115821 | 7040 | 20 | 36 |
| wR6WV | *R. pictipes* | Belem, PA, Brazil | 1127350 | 6027 | 41 | 36 |
| wPIC34WC | *R. pictipes* | Belem, Brazil | 1076284 | 2959 | 19 | 36 |
| wPIC2WA | *R. pictipes* | Para, Brazil | 1031047 | 1288 | 22 | 3 |
| Mean (without wRobQ) |  |  | 1082587±64807 |  |  |  |

The level of read coverage used to distinguish the contigs deriving from the symbionts from those hosted by the genome showed that *Wolbachia* contigs display a 5 to 20-fold higher coverage than contigs from the host genomes (Supplementary Figure 1). Indeed, by re-mapping the reads to the assemblies excluding the incomplete *w*RobQ assembly, we obtained an average coverage of 174X (17X to 370X) for the contigs assigned to *Wolbachia* while the contigs assigned to the host genomes have a medium coverage of 16X (8X to 27X). This result strengthens that these contigs assigned to *Wolbachia* are not the result of lateral gene transfers into host genomes.

In each of the 14 *Wolbachia* assemblies, only one *16S* and one *ftsZ* sequence has been identified. In both cases, these sequences were 100% similar to the sequences obtained previously by PCR. This result suggests that each assembly is composed of sequences belonging to a single *Wolbachia* strain, or alternatively by sequences deriving from a dominant strain.

Global BLASTN searches of the contigs of each assembly against the NR Genbank database revealed a low level of nucleotide divergence (2.6-2.9%) with the *Wolbachia* infecting bedbug, *Cimex lectularius* (*w*Cle). Whole-genome comparisons showed a remarkable level of conservation between *w*Cle and *Rhodnius Wolbachia* (Figure 2).

**Figure 2:**
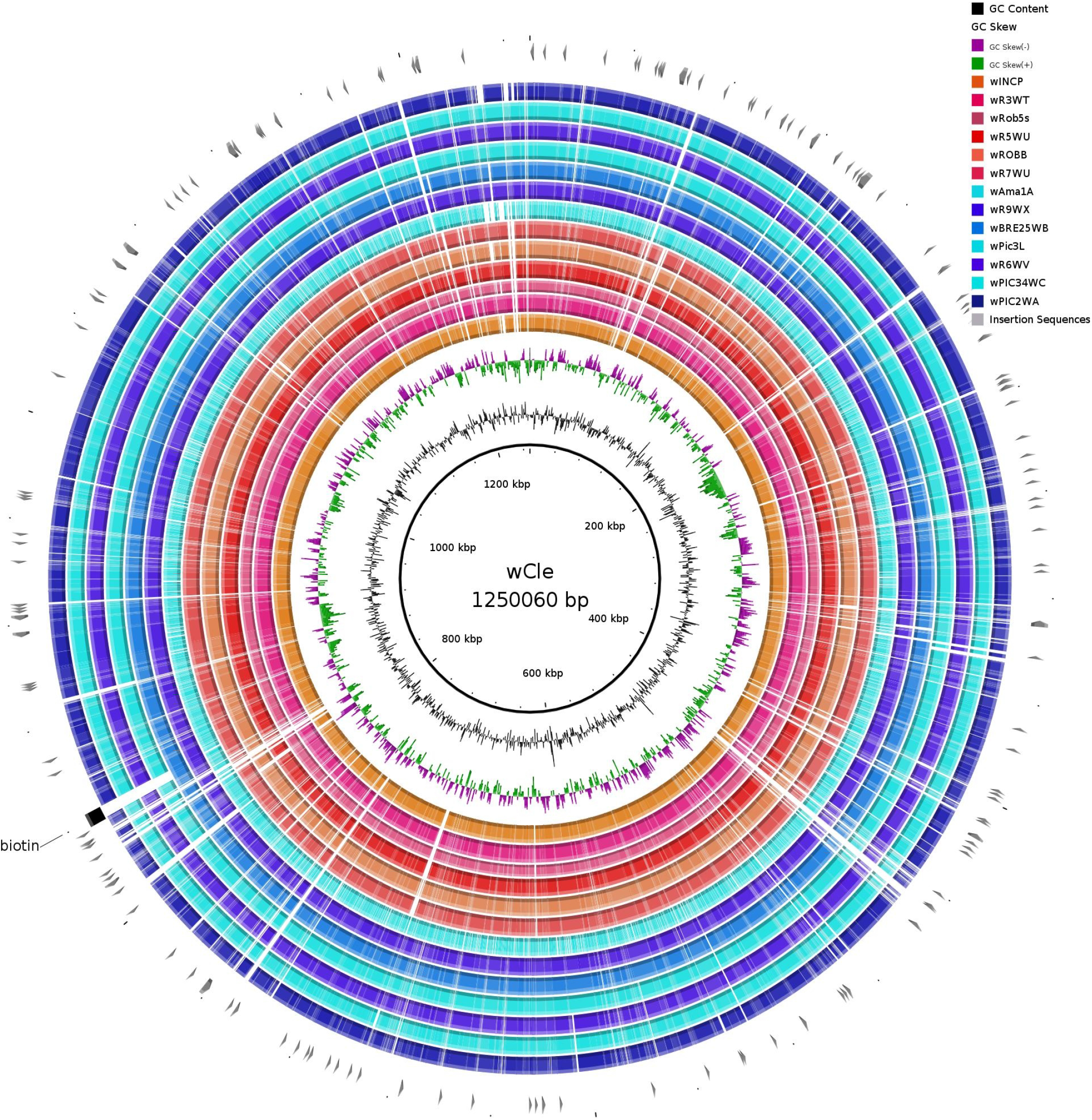
Circular comparison between *Cimex* and *Rhodnius-associated Wolbachia* genomes. Coloured regions correspond to segments with nucleotide similarity (*E*-value < 1e-10, >90% identity). Graphs located in the internal rings indicate GC content and GC skew plots. Blue rings indicate *Wolbachia* genomes infecting the *pictipes* group, whereas orange/red rings represent those infecting the *prolixus* group. In the outer rings, the position of the biotin operons is indicated by a black square and IS-like transposons are represented by grey arrows.

Some gaps correspond to Insertion Sequence (IS) movements, a class of prokaryotic transposable elements that appears to be abundant in these genomes (7-10% of the total genomic content). Interestingly, we can also evidence a deletion at coordinates 835000-840000 shared by *Wolbachia* infecting all the four *R. pictipes* specimens originated from diverse regions. This gap corresponds to one of the rare deletions of a coding region, in this case the one corresponding to the biotin operon (see next section).

We provided a well resolved *Wolbachia* phylogeny based on a supermatrix of 80 conserved genes (Figure 3). This tree confirms the close phylogenetic relationship between *w*Cle and *Rhodnius Wolbachia* in the supergroup F. The phylogenetic positioning of the different *Rhodnius Wolbachia* genomes was globally congruent with the *Rhodnius* tree with the presence of the two major groups, *prolixus* and *pictipes*, supporting the presence of *Wolbachia* in the last common ancestor of the *Rhodnius* lineage and subsequent co-diversification. However, the phylogenetic position of *w*RobQ outside the *prolixus* group is unexpected. Thus, we cannot rule out a whole *Wolbachia* lateral transfer/replacement between members of the different groups.

**Figure 3:**
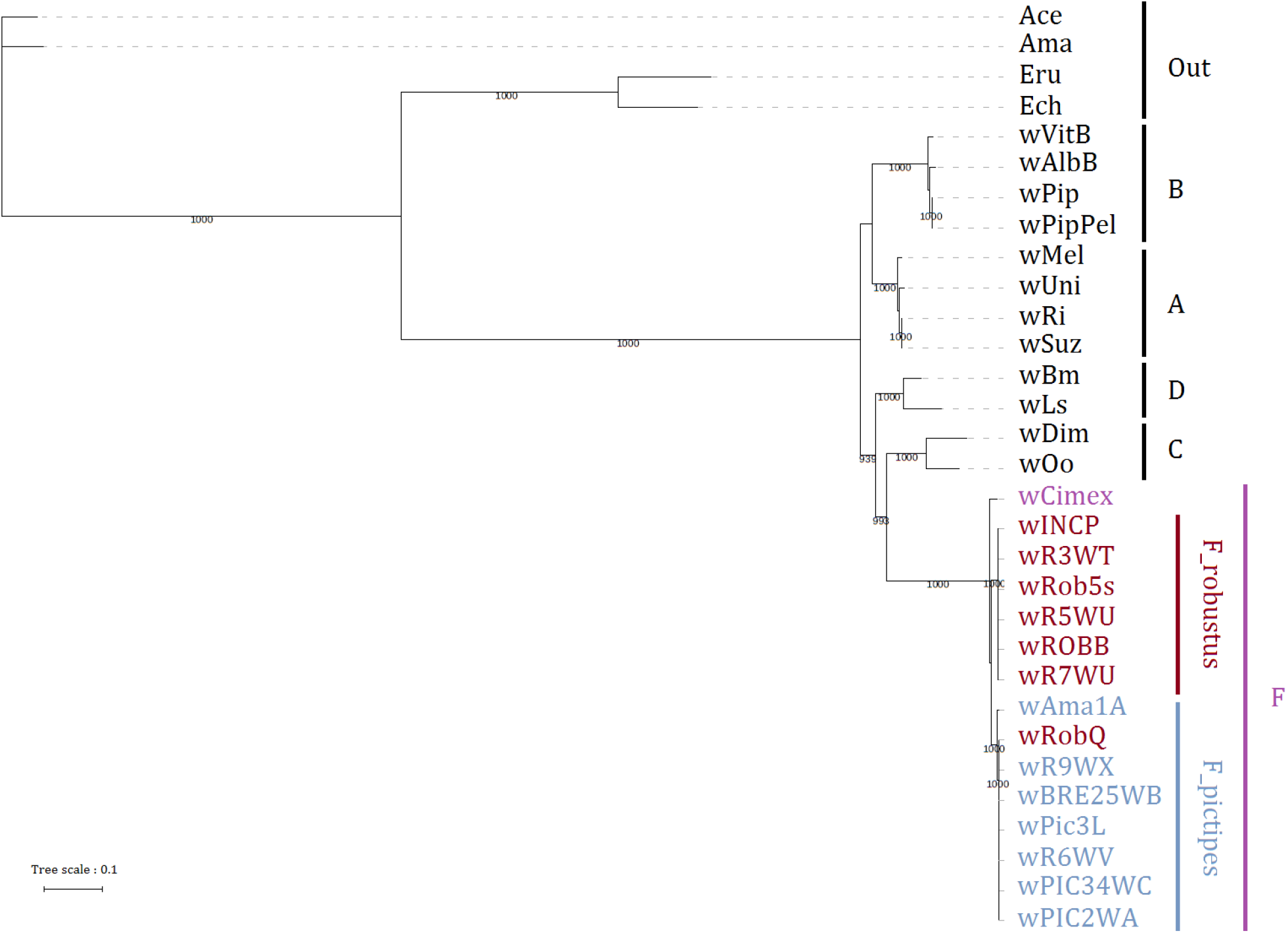
Phylogeny of the different *Wolbachia* strains associated with *Rhodnius* triatomine. The tree represents the best maximum-likelihood phylogeny based on 80 conserved single-copy orthologs. Numbers below nodes indicate the support values (1000 replicates). The dataset used was retrieved from Comandatore *et al*., 2013 for: *Wolbachia* endosymbiont from: *Drosophila melanogaster, w*Mel; *D. simulans, w*Ri; *D. suzukii, w*Suz; *Muscidifurax uniraptor, w*Uni; *Culex quinquefasciatus* JHB, *w*Pip; *C. quinquefasciatus* Pel, *w*Pip Pel; *Nasonia vitripennis, w*VitB; *Aedes albopictus*, wAlbB; *Brugia malayi, w*Bm; *Onchocerca ochengi, w*Oo; *Dirofilaria immitis, w*Di*; Anaplasma centrale* str. Israel, Ace; *Anaplasma marginale* str. Florida, Ama; *Ehrlichia chaffeensis* str. Arkansas, Ech; *Ehrlichia ruminantium* str. Gardel, Eru; *Cimex lectularius, wCimex*. All the *Wolbachia* symbionts from *Rhodnius ssp*. were obtained in this study (see Table 1 for sample denomination). Letters A, B, C, D, F indicate *Wolbachia* supergroups.

### Analysis of the B vitamin genes among *Rhodnius Wolbachia* and *Rhodococcus* strains

As *w*Cle is known to contribute to the fitness of its host by provisioning biotin and riboflavin, we thus analysed the level of conservation of the different B vitamin pathways in the 13 *Rhodnius Wolbachia* genomes (Figure 4a). Moreover, four complete genomes of the triatomine gut symbionts available on the NCBI databank, belonging to the genera *Rhodococcus* and associated with *R. prolixus* and *Triatoma sp*., were also used.

**Figure 4:**
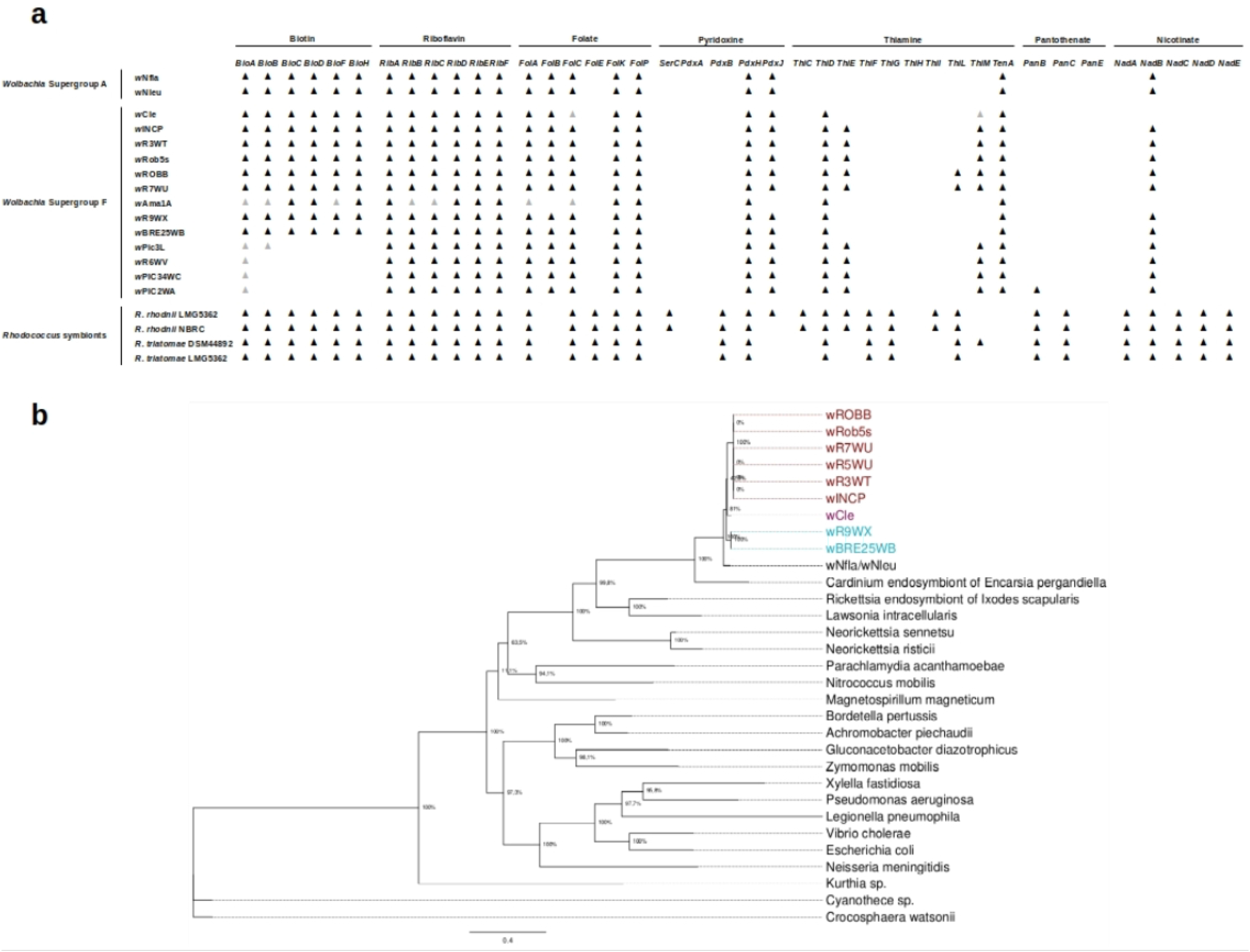
Distribution and evolution of the B-vitamin biosynthetic pathways in *Wolbachia* and *Rhodococcus* symbionts. **(a)** Presence/absence of biosynthetic pathways for B vitamins among *Wolbachia* and *Rhodococcus* symbiont genomes. Black triangles indicate the presence of apparently intact genes, whereas grey triangles represent pseudo-genes. **(b)** Maximum-likelihood phylogeny of the biotin operon. The tree results from the concatenation of the 6 biotin genes. Taxa in brown represent *Wolbachia* infecting the *prolixus* group, in blue, the *pictipes* group and in magenta for wCle. Percentages at nodes indicate the bootstrap supports and the scale bar represents the average number of substitutions per site.

Our analysis showed that the *Rhodococcus* symbionts encode complete biotin, riboflavin and nicotinate operons and nearly complete folate and pantothenate pathways. On average, the triatomine-associated *Rhodococcus* genomes encode for 33 B-vitamin genes (maximum 35), whereas *w*Cle and *Rhodnius Wolbachia* encode for 21 (maximum 24).

Among the B-vitamin pathways only the riboflavin genes are conserved among the group F *Wolbachia*. A complete biotin operon is present in 11 group F *Wolbachia* genomes. Biotin operon is a very rare attribute in *Wolbachia* genomes: our searches in generalist sequence databases using similarity-based analysis have revealed that intact operons are only present in *w*Cle, in two bee-associated *Wolbachia* (*w*Nfla and *w*Nleu) and in *Rhodnius Wolbachia*. In the *Rhodnius Wolbachia*, biotin operon appeared intact in 8 out of the 13 genomes examined, with coverage levels comparable to those of the other parts the *Wolbachia* genomes (Table 3). In the *w*Am1a genome, the biotin operon was fragmented and displays some deletions; moreover, the read coverage of the operon was two times less than the average coverage of the genome. It is thus possible that this pattern results from an ongoing process of biotin operon disruption, but we cannot rule out some bias linked to the incompleteness of the genome assembly as we observed the absence of the biotin genes in wROBQ incomplete genome. However, we observed a deletion of the biotin operon in the *Wolbachia* genome associated with all the four *R. pictipes* specimens (*w*Pic3L, *w*R6WV, *w*PIC34WC, *w*PIC2WA) for which the genome size seems complete (> 1 Mb). Compared to the read coverage of the corresponding *Wolbachia* genomes, only a low proportion of reads matched the biotin operon (Table 3).

**Table 3:** Evolution of the biotin operon. Analysis of the read coverage and the selective pressures (Ka/Ks) acting on the biotin genes in the genomes of the *Wolbachia* F supergroup. NA: Not Applicable.

| *Wolbachia* | Read coverage (X) |  | Ka/Ks |  |  |  |  |  |
| --- | --- | --- | --- | --- | --- | --- | --- | --- |
|  | Biotin | Genome | BioA | BioB | BioC | BioD | BioF | BioH |
| wCle | NA | NA | 0.139741 | 0.0518431 | 0.246587 | 0.0769836 | 0.0947696 | 0.0703239 |
| wBRE25WB | 46 | 56 | 0.138603 | 0.049762 | 0.273816 | 0.0606412 | 0.0898628 | 0.0754634 |
| wINCP | 17 | 17 | 0.122456 | 0.05335 | 0.296629 | 0.0706004 | 0.0846851 | 0.0675378 |
| wROBB | 45 | 46 | 0.122456 | 0.05335 | 0.296629 | 0.0706004 | 0.0846851 | 0.0675378 |
| wRob5s | 44 | 141 | 0.122456 | 0.05335 | 0.296629 | 0.0706004 | 0.0846851 | 0.0675378 |
| wR9WX | 28 | 32 | 0.138603 | 0.049762 | 0.273816 | 0.0606412 | 0.0898628 | 0.0754634 |
| wR3WT | 326 | 370 | 0.122456 | 0.05335 | 0.296629 | 0.0706004 | 0.0846851 | 0.0675378 |
| wR5WU | 228 | 265 | 0.122456 | 0.05335 | 0.296629 | 0.0706004 | 0.0846851 | 0.0675378 |
| wR7WU | 250 | 327 | 0.122456 | 0.05335 | 0.296629 | 0.0706004 | 0.0846851 | 0.0675378 |
| wAma1a | 37 | 68 | disrupted | disrupted | 0.308917 | 0.0731896 | disrupted | 0.0710802 |
| wPic3L | 39 | 244 | disrupted | disrupted | absent | absent | absent | absent |
| wR6WV | 24 | 257 | disrupted | absent | absent | absent | absent | absent |
| wPIC34WC | 19 | 226 | disrupted | absent | absent | absent | absent | absent |
| wPIC2WA | 19 | 213 | disrupted | absent | absent | absent | absent | absent |

Using local alignments of the contigs encoding the *Rhodnius Wolbachia* biotin operons, it was possible to identify the location of the deletion (Supplementary Figure 2). This deletion was located at exactly the same position in the four genomes: in the middle of the bioA gene and at the end of the bioB gene. These data allowed us to exclude potential assembly artefacts and suggested that a single deletion arose in the ancestors of these four *Wolbachia* strains rather than occurrences of independent and multiple deletions exactly at the same genomic coordinates. Finally, the genes involved in the other B-vitamin pathways displayed an erratic distribution in supergroup F *Wolbachia* except for the riboflavin operon that appeared well conserved (Figure 2).

Concatenated phylogeny of the 6 biotin genes composing the operon indicated that *Rhodnius Wolbachia* and *w*Cle form a well-supported monophyletic group (Fig 4b). The individual phylogenies of the biotin genes also gave the same topology (Supplementary Figure 3).

Finally, we estimated the ratio of non-synonymous versus synonymous substitution rates (Ka/Ks) in the biotin genes, a proxy of the selective constraints acting on these genes (Table 3). The Ka/Ks ratio is comparable and below 1 in *w*Cle and *Rhodnius Wolbachia*, indicating strong purifying selection pressures.

### Pervasive lateral gene transfers between *Rhodnius Wolbachia* and *Rhodnius* genomes

*Wolbachia*-to-eukarya lateral gene transfers (LGTs) constitute interesting proof-prints of past infections. We examined the 36 *Rhodnius* genomes for *Rhodnius Wolbachia* genes. A gene with >90% of similarity with a *Rhodnius Wolbachia* gene was considered to be integrated in the host genome only if its 3’ and the 5’ boundaries aligned on the *R. prolixus* reference genomes, providing a proof that these *Rhodnius Wolbachia*-originated genes were embedded in a *Rhodnius-like* context. Using these criteria, Figure 5 displays for the 36 *Rhodnius* samples the corresponding location of these putative LGTs mapped in the *w*Cle genome. This allowed us to visualize the extent and the diversity of the *Rhodnius Wolbachia* genes integrated into the host genomes. With the exception of the *pallescens* group, most of the *Rhodnius* genomes displayed many *Rhodnius Wolbachia* genes, including samples in which no *Wolbachia* infection was evidenced by PCR. For some genomes, the amount of gene transfers was significant (Figure 6), up to 350kb for *R. sp*. INCP (>240 segments). In fact, almost all of the *Rhodnius Wolbachia* genes have been transferred into the host genomes at some point. Re-mapping of the reads on the laterally transferred genes indicated that the level of coverage was very close to the medium coverage of the host genome.

**Figure 5:**
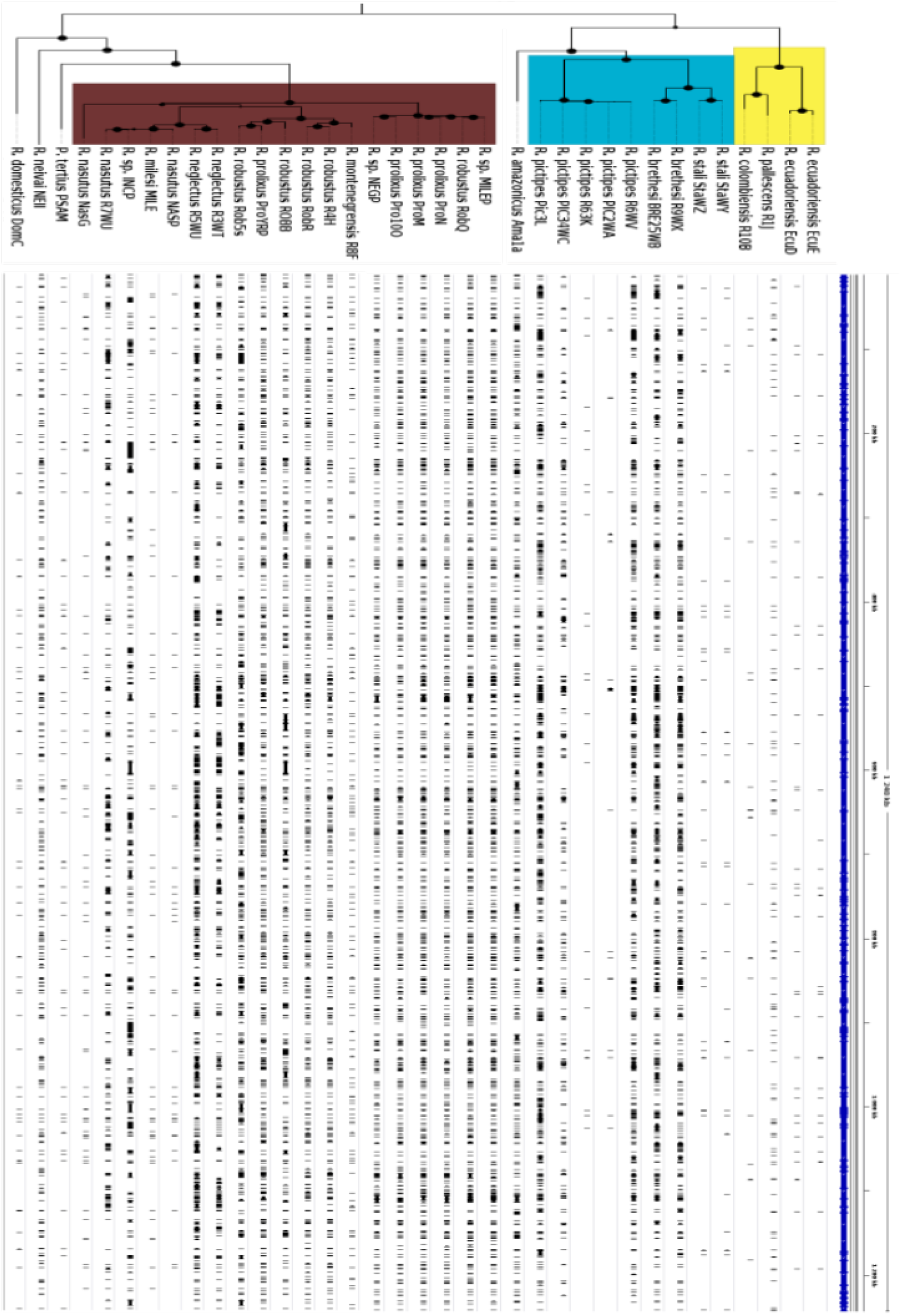
Evidence of laterally-acquired *Wolbachia* genes in the *Rhodnius* genomes. Mapping of the *Wolbachia* genes integrated the host genomes against the *w*Cle genome. Each black segment represents sequences in the corresponding *Rhodnius* genomes and aligned against the *w*Cle genomes with >95% of nucleotide similarity and *E*-value < 1e-10. The tree represents the *Rhodnius* mitochondrial genome maximumlikelihood phylogeny. Yellow, blue and brown groups in the tree refer to the *pallescens, pictipes* and *prolixus* groups respectively. Black circles in the phylogeny indicate the support values of each node: large circles for bootstraps >99%, small ones for supports between 90% and 99%.

**Figure 6:**
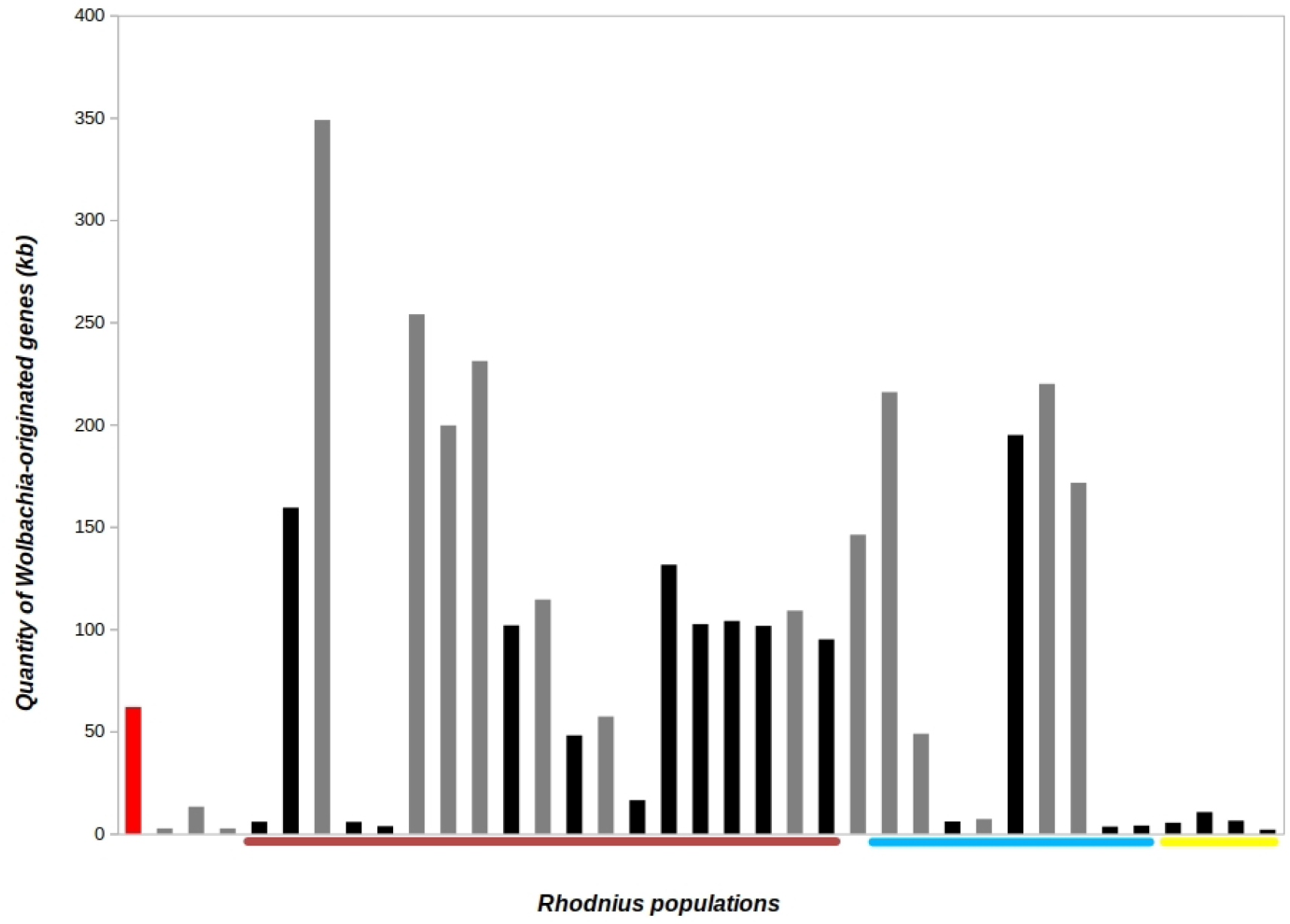
Amount of *Wolbachia* genes in the *Rhodnius* genomes. The red bar indicates the amount of transferred DNA in the *R. prolixus* reference genome, whereas the grey and black bars represent *Rhodnius* samples in which *Wolbachia* infection have been and have not been evidenced respectively. Yellow, blue and brown lines below the plot refer to the *Rhodnius* genomes belonging to the *pallescens, pictipes* and *prolixus* groups respectively in the same order as in the phylogeny in the Figure 5.

For example, in the *Wolbachia*-free ProN assembly, the global level of coverage of the laterally transferred genes was 19x, similar to the global genome coverage (15x). These results strongly suggest that the possible cases of LGTs identified here correspond to integrated genes and are not the result of false assignments between DNA segments present in the *Wolbachia* genomes and those present in the *Rhodnius* genomes. It is worth noting that integrated *Wolbachia* genes are overrepresented at the boundaries of the *Rhodnius* contigs. Indeed, 30% of them are located at less than 500nt from the contig ends (Supplementary Figure 4). Thus, we can not ruled out that some of these *Wolbachia* insertions located at the boundaries of the host genome contigs correspond to false chimeric sequences. Finally, the presence of *Rhodnius Wolbachia* genes in nearly all the *Rhodnius* genomes tested in this study indicates that the level of infection by *Wolbachia* was widespread and global in the genus. Even in samples in which *Wolbachia* associations have not been evidenced by PCR, genetic traces of past infections remain. For comparison, we analysed the published *Rhodnius prolixus* reference genome for the presence of *Rhodnius Wolbachia* genes. We have evidence for 180 genes or gene fragments resulting from LGTs that are scattered on 45 contigs.

## Discussion

Blood-sucking insects are often associated with symbiotic bacteria for provisioning some nutrients that are naturally deficient in the blood of their hosts. This phenomenon has been called “nutritional symbiosis”. For triatomines of the *Rhodnius* genus, it was assumed that the gut bacterial symbionts *R. rhodnii* play the role of nutritional symbionts by providing their hosts with B vitamins (Hill et al. 1976). Here, we report that the situation might be more complex because many *Rhodnius* populations also display associations with a group of *Wolbachia* that could be nutritional symbionts in insects (Hosokawa et al. 2010) and in nematodes (Keiser et al. 2008). Using a combination of PCR, whole-genome sequencing and diverse genetic and genomics analyses conducted on a large set of *Rhodnius* species and populations, we tried to address the origin, nature, ecological and evolutionary meanings of this association.

### *Rhodococcus rhodnii*, *Wolbachia* and *Rhodnius* compose a widespread and probably ancient association

Comparable genome sizes for *Wolbachia* from *Rhodnius* to published complete *Wolbachia* genomes were found, i. e. around 0.7-1.5Mb (Lindsey et al. 2016, Manoj et al. 2021). Larger genomes are described for *Rhodoccocus*, namely for *R. rhodnii* and *R. triatomae* from 4.38 to 5.8 Mb. The differentiated size of these two bacteria genomes reflects their lifestyle: *Wolbachia* are ancient parasites, with highly reduced/degraded genomes (Gerth et al. 2014), whereas *Rhodococcus* belong to a very diverse group of environmental, free-living bacteria (rarely pathogenic and symbiotic) with large genomes (Bell et al. 1998). We have shown that the phyletic distribution of the gut symbiont *R. rhodnii* and the *Wolbachia* infecting *Rhodnius* bugs is different: whereas *R. rhodnii* was detected in all the triatomine samples tested, *Wolbachia* displays a patchy distribution, infecting 40% of the samples (corresponding to eight *Rhodnius* species out a total of 17). The infection was not species-dependant, as for a given species some strains were infected and others were not (as for *R. pallescens*, for example). Moreover, one *R. pictipes* lab strain was *Wolbachia-free* out of the five tested, and the absence of *Wolbachia* was confirmed using whole-genome sequencing. However, the analysis of *Wolbachia*-to-host gene transfers indicated that almost *Rhodnius* species from *prolixus* and *pictipes* group harbour *Rhodnius Wolbachia*-originated genes, including samples in which present *Wolbachia* infection was not evidenced by PCR or by genome high throughput sequencing. We found at least four *coxA* and one *ftsZ* genes inserted into the host chromosomes. In some extent, these nuclear insertions might also bias PCR survey for estimating *Wolbachia* prevalence due to the amplification of host nuclear *Wolbachia* genes rather than those coming from free-living *Wolbachia*. Moreover, false positive PCR could be due to some *Wolbachia* contamination by parasites harbouring such bacteria as nematodes. In our data, no (of very few) contigs in our *Rhodnius* assemblies match with nematode genes (data not shown). Moreover, the integrated *Wolbachia* genes into *Rhodnius* genomes, as observed in the *Brugia malayi* genome (Ioannidis et al. 2013), are over-represented at the boundaries of the *Rhodnius* contigs. Many cases of acquisition of *Wolbachia* genes by their host have been reported in the literature (Dunning Hotopp 2011), and the analysis of the reference *R. prolixus* genome has revealed the presence of 25 putative cases of LGTs (Mesquita et al. 2015). Using a larger collection of *Rhodnius* genomes and the corresponding *Wolbachia* genome sequences, we showed that the events of gene transfers have been considerably more frequent and massive than initially suspected. Interestingly, these events also provide molecular signatures of past infection, which could be used to infer the prevalence of the *Wolbachia* infection (Koutsovoulos et al. 2014; Keroack et al. 2016). Our results indicate that almost all the *Rhodnius* species have been infected by *Wolbachia*, supporting the view that the *R. rhodnii, Wolbachia* and *Rhodnius* association is a widely distributed and probably an ancient phenomenon, preceding the diversification of the genus *Rhodnius*. Whereas the *R. rhodnii* symbiosis composes a stable association, *Rhodnius Wolbachia* co-infection appears to be a more dynamic process with events of recurrent losses and gains. Co-symbiosis with several bacterial partners has been recently evidenced in many insect lineages, especially in hemipteran insects (Sudakaran et al. 2017). Sometimes, competition and replacement occur between the different microbes. However, in many cases, dual symbiosis is also present (Sudakaran et al. 2017). If there is little doubt that *R. rhodnii* acts as a mutualist symbiont with the triatomine by supplementing the host blood diet with B vitamins (Hill et al. 1976), the evolutionary origins and the true nature of the relationship between *Rhodnius* and *Wolbachia* is even more puzzling.

### The close relationship between *Rhodnius Wolbachia* and *Wolbachia*-infecting bedbugs support one or more host switches

To better document the origin and the nature of the relation between *Rhodnius Wolbachia* and triatomine bugs, we assembled and analysed 13 *Rhodnius Wolbachia* genomes. *Rhodnius Wolbachia* genomes are highly similar to the genome of the *Wolbachia* that infects the bedbug *Cimex lectularius* (*w*Cle). Indeed, *Rhodnius Wolbachia* and *w*Cle display a very high level of genome conservation, gene similarities and phylogenetic affinities, which strongly suggests that these *Wolbachia* share a very recent common ancestor. This result is unexpected as bedbugs and triatomines are distantly related Hemipteran insects with a divergence time since their last common ancestor estimated at around 185My (Hwang and Weirauch 2012). Our data contradict the possibility of a vertical inheritance of *Rhodnius Wolbachia* and *w*Cle since their last common ancestor and favour a scenario in which host switches and lateral acquisition have occurred. Based on multi-locus sequence phylogeny or trans-infection experiments in the laboratory, several studies have documented the existence of lateral acquisitions of *Wolbachia* between distantly related species (Vavre et al. 1999; Ahmed et al. 2015; Ahmed et al. 2016; Lefoulon et al. 2016). Our study provides evidence at the genome level that *Wolbachia* host switches, followed by a long-term establishment in nature, are not associated with major genome recombination. Indeed, with the exception of a few insertions/deletions sometimes associated with Insertion Sequence movements, *Rhodnius Wolbachia* and *w*Cle genomes appear highly stable and cohesive, suggesting that host switches are not a steep slope that requires, or generates, major genomic changes. If *Rhodnius Wolbachia* and *w*Cle have undergone a relatively recent host switch, given their extremely high level of overall genome similarities, the direction and origin of the host transfers remain speculative. These *Wolbachia* belong to the F supergroup, one of the less known clusters of the family, and compose a heterogeneous assemblage of microbes infecting diverse arthropods and nematodes (Ros et al. 2009, Manoj et al. 2021). Although the paucity of the genetic and genomic data concerning the F supergroup precludes any conclusion regarding the origin and evolution of this lineage, a direct *Wolbachia* transfer between an ancestor of bedbugs and an ancestor of the *Rhodnius* triatomine could be suggested. Alternatively, a direct host switch between *Cimex* and *Rhodnius* may have occurred. Many Cimicidae species present in South America feed on bats and birds, as do the triatomine species (Poggio et al. 2009; Georgieva et al. 2017). The prevalence and the nature of the *Wolbachia* infections of these cimicids are unknown, but close contacts, interactions and possible microbe exchanges between them and triatomines appear plausible, since cannibalism and coprophagy have been described in Triatominae (Schaub et al. 1989). Interspecific haemolymphagy and cleptohaematophagy are demonstrated for some triatomines, which may be an extra source for exchanging the microbiota (Durán et al. 2016). But two independent transfers from a third player cannot be ruled out. For example, parasitic nematode that are known to harbour F *Wolbachia* strain, could be such candidate. Additional data on the distribution and the nature of the *Wolbachia* in the Cimicidae and Triatominae families would bring valuable information to resolve the close relationship between *Rhodnius Wolbachia* and *Wolbachia*-infecting bedbugs.

### Do *Wolbachia* maintain a mutualistic relationship with *Rhodnius?*

Even with the lack of functional or *in vivo* experimentations to document the fitness advantages potentially provided by *Wolbachia*, the very close phylogenetic relatedness and the remarkable genomic similarities between *w*Cle and *Rhodnius Wolbachia* open the possibility that the nutritional mutualism in *Rhodnius* might be carried out by other symbionts than *Rhodococcus*. Mutualism in supergroup F *Wolbachia* has been documented in the bedbug *Cimex lectularius* (Hosokawa et al. 2010) but has also been suggested in the nematode *Mansonella perstans* (Keiser et al. 2008), indicating that mutualism may be common in this supergroup (Gerth et al. 2014). The presence of a biotin operon in the genome of *w*Cle has been identified as the key determinant of nutritional mutualism based on the B-vitamin supplementation of the host blood diet and it was assumed that the presence of this operon in *w*Cle was the result of a lateral gene transfer from an unidentified cosymbiont (Nikoh et al. 2014). While for the B-vitamin pathways the riboflavin genes are conserved among the group F *Wolbachia*, reflecting the general situation in the *Wolbachia* genus (Moriyama et al. 2015), the occurrence of a biotin operon in *Wolbachia* genomes is very rare (Gerth and Bleidorn 2016). However, a highly disrupted and mutated biotin operon has been identified in *Wolbachia* infecting the nematode *Onchocerca ochengi (wOo*)(Nikoh et al. 2014). But it is worth noting that for some obligate blood feeders the transfer capabilities of the biotin operon are not limited to *Wolbachia* (Duron & Gottlieb, 2020) since diverse B vitamin-provisioning symbionts harbouring related biotin operons have been detected including *Midichloria* and *Rickettsia* in the invasive tick *Hyalomma marginatum* (Buysse et al., 2021) and *Legionella* in the louse *Polyplax serrata* (Říhová et al., 2017). In our study, we documented the presence of a biotin operon in *Rhodnius Wolbachia* genomes for at least 8 genomes infecting two *Rhodnius* groups, *prolixus* and *pictipes*. The operon is intact and under strong selective constraints, comparable to those acting on the *w*Cle genome. This result suggests that the biotin operon in these 8 genomes is functional and might contribute to host fitness. As observed in wOo, a deletion of the biotin operon has occurred in 4 *Wolbachia*-infecting *R. pictipes* hosts, suggesting a possible breakdown of the nutritional symbiosis. The phylogeny of the biotin genes indicates an existing presence of the operon in the ancestors of *w*Cle and *Rhodnius Wolbachia*, suggesting that they have been stably maintained in most *Rhodnius Wolbachia* strains over time. Taken together, these results suggest that the biotin operons in *Rhodnius Wolbachia*, along with other well conserved B-vitamin operons, such as riboflavin, might be involved in a nutritional symbiotic relationship, as observed with *w*Cle. As the gut symbiont *R. rhodnii* is orally transmitted, whereas *Wolbachia* is transmitted *via* the maternal lineage, it could be selectively advantageous to maintain a symbiotic system in which *Wolbachia* might eventually compensate the transitory absence of the *R. rhodnii* symbionts. Interestingly, bedbugs also harbour secondary endosymbionts belonging to the γ-proteobacteria family (BEV-like symbionts). Whereas *Wolbachia* prevalence is high, BEV-like symbionts are scarcer (Meriweather et al. 2013). However, suppression of BEV-like symbionts by an antibiotic treatment led to a reduction in the fertility of the bedbugs (Sakamoto and Rasgon 2006). Given the close relatedness between *Rhodnius Wolbachia* and *w*Cle, the symmetry of the situation in bedbugs and in triatomine is striking, except that the role of the *Wolbachia* might be inverted: an obligatory symbiont in bedbugs and probably a facultative one in triatomine bugs.

Finally, the possible presence of *Wolbachia* able to synthesize biotin and other B-vitamins might also explain some contradictory results obtained in survival tests of *Rhodnius* larvae cured for *R. rhodnii* (Hill et al. 1976). As the presence of *Wolbachia* was never checked in these experiments conducted between the 1950s and 70s (Baines 1956; Lake and Friend 1968; Nyirady 1973; Auden 1974; Hill et al. 1976), the presence of some strains by *Wolbachia* is plausible.

### Conclusion and perspectives

Triatomine bugs and bedbugs are distantly related species that have in common the need to feed on vertebrate bloods, an adaptation made possible by the presence of symbiotic microbes that supplement their diet with B vitamins. Surprisingly, these bugs also share very closely related *Wolbachia* symbionts that most probably results from lateral exchanges. Our results suggest that *Wolbachia* has the potantial to act as a nutritional mutualist in triatomines, as observed in bedbugs, in complementation (or in rescue) to the *R. rhodnii* gut symbionts. Ultimately, only *in vivo* functional tests with triatomines cured with *R. rhodnii* and/or *Wolbachia* will address the exact symbiotic role of each microbe. Otherwise, genetic manipulation of insect vectors could be a potential alternative to chemical strategies to control vector populations. Paratransgenesis approach, involving genetic manipulation of commensal or symbiotic bacteria harbouring by vector hosts, was applied to *R. rhodnii* (Beard et al., 1993). For example, experimentally, the symbiont has been transformed to export Cecropin A, lethal to the Chagas’disease parasite, *Trypanosoma cruzi* (Durvasula et al., 2003). However, field applications require long-term safety studies. The fact that many *Rhodnius* species are infected with *Wolbachia* also opens up a paratrangenesis alternative since *Wolbachia*-based strategies to control insect pests and disease vectors has been developed (Brelsfoard et al., 2009) and for instance are investigated in controlling *Aedes*-borne viral transmission (Ogunlade et al., 2021). We believe that genomic data analysed in this study shed new light on the origin and the evolution of the nutritional symbiosis in the *Rhodnius* vectors.

## Acknowledgements

The authors thank Céline Chaillot and Shirley Hé for some PCR experiments and Morgane Lavina and Claire Capdevielle-Dulac for technical assistance.

We thank Clement Gilbert for comments on the manuscript and advice in the course of the project, Jean-Michel Rossignol and Carlos Eduardo Almeida for careful reading of the final version of the paper and the two reviewers for their helpful recommendations.

We thank Malcolm Elden for the professional English spelling corrections.

